# High-quality genome assembly of *Cinnamomum burmannii* (chvar. Borneol) provides insights into the natural borneol biosynthesis

**DOI:** 10.1101/2021.09.14.460210

**Authors:** Fangping Li, Shilin Huang, Yu Mei, Bingqi Wu, Zhuangwei Hou, Penglin Zhan, Shike Cai, Qingmei Liu, Zhihao Hou, Junliang Zhao, Jihua Wang

**Affiliations:** Guangdong Provincial Key Laboratory of Crops Genetics & Improvement, Crops Research Institute, Guangdong Academy of Agricultural Sciences, Guangzhou 510640, China; Rice Research Institute & Guangdong Key Laboratory of New Technology in Rice Breeding, Guangdong Academy of Agricultural Sciences, Guangzhou 510640, China; Guangdong Provincial Key Laboratory of Plant Molecular Breeding, State Key Laboratory for Conservation and Utilization of Subtropical Agro-Bioresources, South China Agricultural University, Guangzhou 510642, China; State Key Laboratory of Biocontrol and Guangdong Provincial Key Laboratory of Plant Resources, School of Life Sciences, Sun Yat-sen University, 135, Xingangxi Road, Guangzhou, Guangdong, 510275, China; Guangdong Province Key Laboratory of Microbial Signals and Disease Control, Integrative Microbiology Research Centre, South China Agricultural University, Guangzhou 510642, China; Guangdong Provincial Engineering & Technology Research Center for Conservation and Utilization of the Genuine Southern Medicinal Resources, Guangzhou 510640, China

**Keywords:** *C. burmannii* (chvar. Borneol), Genome, D-borneol

## Abstract

*Cinnamomum burmannii* (chvar. Borneol) is a well-known medicinal and industrial plant cultivated in the Lingnan region of China. It is the key source from organism of natural borneol (D-borneol), one of the precious and widely used Chinese herbal medicines with a variety of medicinal effects. Here, we report a high-quality chromosome-scale genome assembly of *C. burmannii* (chvar. Borneol) using Pacbio single-molecule sequencing and Hi-C technology. The assembled genome size was 1.14 GB with a scaffold N50 of 94.30 Mb, while 98.77% of the assembled sequences were anchored on 12 pseudochromosomes including 41549 protein-coding genes. Genomic evolution analysis revealed *C. burmannii* and *C. micranthum* shared two Lauraceae unique ancestral whole-genome duplication (WGD) events. Likewise, comparative genomic analysis showed strong collinearity between these two species. Besides, the analysis for Long Terminal Repeat Retrotransposons (LTR-RTs) indicated the outbreak of LTR-RTs insertion made a great contribution to the size difference of genomes between *C. burmannii* and *C. micranthum.* Furthermore, the candidate genes in pathway associated with natural borneol synthesis were identified on the genome and their differential expressions were analyzed in various biological tissues. We considered that several of genes in Mevalonate (MVA) Methylerythritol Phosphate (MEP) pathways or in downstream pathway have the potential to be the key factors in the biosynthesis of D-borneol. We also constructed the genome database (CAMD; http://www.cinnamomumdatabase.com/) of *Cinnamomum* species for a better data utilization in the future. All these results will enrich the genomic data of Lauraceae plants and facilitate genetic improvement of this commercially important plant.

## Introduction

Natural borneol, also known as D-borneol, borneol, plum borneol, plum flower, etc., is a highly desirable natural product, as well as a precious Chinese herbal medicine. It has been widely used as folk medicine and food additives in China and India (Chen et al., 2010). Natural borneol has the effect of opening the orifice and awakening the mind, clearing away heat and relieving pain. It is often utilized for the treatment of multiple diseases such as sore throat, convulsive phlegm and shiver. Previous researches indicated D-borneol has remarkable effects on the aspects of anti-myocardial ischemia, brain tissue protection, nerve protection and anti-infection *in vivo* or *in vitro* (Armaka et al., 1999; Lu et al., 2010). Natural borneol is also a fragrance ingredient used in decorative cosmetics products. Its use worldwide is in the region of 1–10 metric tones per annum (Bhatia et al., 2008). As of 2016, the annual demand for natural borneol in the world was about 12,000 tons, and the total annual output of natural borneol in China was less than 200 tons, indicating a huge market gap and potential (http://www.ibaogao.com/baogao/10241a2312016.html). As an important source of natural borneol, *Cinnamomum burmannii* (chvar. Borneol) also called is a well-known medicinal and edible plant which had been widely cultivated in the Lingnan region of southern China (Wolf et al., 2011). Phytochemical studies have shown that *C. burmannii* (chvar. Borneol) contains abundant natural borneol in its leaves (Huang et al., 2019). In China, Guangdong Province is the main planting area for *C. burmannii* (chvar. Borneol), accounting for approximately 90% of its output (Chen et al., 2010).

Based on the identified metabolites and enzymes, previous studies have determined that the biosynthesis pathway of Glycerol 3-Phosphatase (GPP), the precusor of D-borneol *in vivo,* is related to the MVA pathway. Recently, MEP pathway in plants also had been reported to be associated with the GPP formation. Previous transcriptomic and synthetic biology studies also had identified that a gene (CbTPS) in TPS gene family is a key factor for downstream transformation of GPP generated by upstream pathways and natural D-borneol in *C. burmannii* (Ma et al., 2021). However, due to the lack of a complete genome, systematic researches of D-borneol biosynthesis and evolutiology of *C. burmannii* at the genomic level were still absent.

In order to fill this gap, we *de novo* assembled a chromosome-scale genome of Mei Pian tree, a physiological type of *C. burmannii* (chvar. Borneol) and whose leaves is rich in D-borneol. The assembly was based on single-molecule real-time long reads from the Pacific Biosciences (PacBio) SequeII platform, and construct pseudo-chromosomes using high-throughput chromosome conformation capture (Hi-C) techniques base on a chromosome number of 2n = 24 as previous report (Wu et al., 1992).

By evolution and comparative genomic analysis, we detected that the recent insertion outbreak of LTR-RTs was an important cause of plant differentiation in *Cinnamomum*. RNA-seq transcriptiomics and further gene annotation analysis based on our assembled *C. burmannii* genome dissected the natural D-borneol biosynthesis pathway at the level of genome. The present study also demonstrated, *CbERG20* and *CbIDI1* in MVA pathway, *CbDXR*, *CbCMS* and *CbHDR* in MEP pathway as well as *CbTPS-1* in the downstream pathway, tend to be the key or the limiting factor in the natural D-borneol biosynthesis pathway in *Cinnamomum*. To encourage the data utilization, we also built a user-friendly genomic database of *Cinnamomum* (CAMD; http://www.cinnamomumdatabase.com/) for the data access.

The present study provides the first reference genome and detail genomic information for *Cinnamomum burmannii* (chvar. Borneol), a plant rich in D-borneol which is highly desired natural product used in medicine and industry. We also dissected the D-borneol synthesis pathway and provide a novel insight into the molecular mechanisms of this pathway. Our results will be valuable resources for promoting genetic improvement and understanding the biosynthesis of active ingredients of *C. burmannii* (chvar. Borneol).

## Method and materials

### Plant materials and sequencing

Root, leaves, and stem samples of “Mei 3”, a popular cultivated variety of *Cinnamomum burmannii* (chvar. Borneol) were collected from the Mei Zhou City (Pingyuan District, 24°28’–24°66’ N, 117.3°13’–118.1°21’ E) in Guangdong Province, China. Because of its unique-geography and environment, the Pingyuan District of Mei Zhou City is considered the authentic production area of *C. burmannii*. “Mei 3”, with many excellent cultivation characteristics, including growth fast, high quality of component, and disease resistance, is the main variety of *C. burmannii* (chvar. Borneol). In Pingyuan District. High-quality genomic DNA was extracted using a QIAGEN® Genomic Kit (QIAGEN, Germany). The integrity of the DNA was checked by 0.75% agarose gel electrophoresis.

### Genomic survey and assembly

The genommic size and heterozygosis were estimated by the software GCE, Jellyfish, kmergenie and Genomescope (Vurture et al., 2017). The data generated form Pacbio SeqII was implemented to *De novo* assemble with the software MECAT2 (Xiao et al., 2017) and polished by the software NextPolish (Hu et al., 2019) combining the illumina short read data mentioned in survey.

### Hi-C experiment and chromosome assembly construction

To further anchor the contigs to chromosomes, fresh young leaves was utilized to construct a Hi-C library using a NEBNext Ultra II

DNA Library Prep Kit with the enzyme HandIII. 78.71 GB of clean data was generated for Hi-C analyses by the Illumina NovaSeq 6000 platform. For the anchoring of contigs, the mounting process was conducted by the pipeline Juicer (Durand et al., 2016) and 3D-DNA (Dudchenko et al., 2017). Software Juicerbox (Durand et al., 2016) was implemented to visualization and adjustment.

### Genomic assembly quality assessment

Multiple methods were implemented to assess the accuracy and completeness of the assembled genome. First, the paired-end reads were mapped to the genome to evaluate its completeness using BWA-MEM with the default parameters (Li, 2013). RNA-seq data from different tissues (leaves, stalk, and root) were also aligned to the reference genome to obtain the mapping rate using HISAT2 with the default settings (Kim et al., 2015). Second, GC depth scatter plots were used to evaluate any contamination in the sequencing data. Finally, the accuracy and completeness of the genome assembly were evaluated by using BUSCOs to identify the single-copy genes in the assembled genome with the Embryophyta_odb10 database (Simão et al., 2015).

### Gene prediction, Repeat element identification

The sequence of repeat element in *C. burmannii* genome was *de novo* identified by software Repeatmodeler (Flynn et al., 2020) and masked by Repeatmasker (Chen, 2004). The detection and taxonomy about the LTR-RTs was accomplished by LTR_Finder (Xu and Wang, 2007), LTR_harvest (Ellinghaus et al., 2008) and LTR_retriever (Ou and Jiang, 2018).

To generate gene annotation of *C. burmannii* genome, software Augustus was utilized in *de novo* gene prediction. A total of 65.74 GB of transcriptome data from roots, stems, and leaves were alignment to the *C. burmannii* genome by Hisat2 and reassembled into transcripts by Stringties (Pertea et al., 2015) and Transdecoder (Fangping et al., 2021; Kim et al., 2015), which generated the site information of transcriptome. The protein sequences of *C. micranthum* previously published (Chaw et al., 2019) were utilized for the homologous gene prediction by GeMoMa (Keilwagen et al., 2019). The information of *de novo* gene prediction, transcriptome and homologous gene prediction was combining by the software EVM (Haas et al., 2008). PASA and the transcripts generated by Transdecoder in previous step were utilized to update the gff3 file for three rounds to add alternatively spliced isoforms, adjusting the gene structure and add UTR regions to gene models (Haas et al., 2008).

### The Annotation of Gene function

Functional annotation was achieved by using NCBI BLAST + v 2.2.2854 (Altschul et al., 1990) with cutoff e-values of 1e-5 and max target sequences 20 to compare predicted proteins against public databases, including SwissProt, Nr (Plant). The Best-hit BLAST results were then considered as the gene functions. Gene Ontology (GO) identifiers for each gene were obtained using InterProScan-5.25-64.0 by identify motifs and domains by matching against public databases (Zdobnov and Apweiler, 2001). The predicted proteins were also submitted to KAAS to get KO numbers for KEGG pathway annotation (M and S, 2000).

### Phylogenetic analysis and Estimation of divergence time

MAFFT v.7.271 (Katoh and Standley, 2013) with option -maxiterate 1000 was used to align 10 sets of Coding DNA Sequence (CDS) of 230 single-copy orthologous groups. Each orthologous group alignment was used to compute a maximum likelihood phylogeny using iqtree with the nucleic acid substitution model GTR+F+R4 and 1000 bootstrap replicates (Nguyen et al., 2015). The divergence time of each tree node was inferred using MCMCtree of the PAML (Yang, 2007). The final species tree with *Amborella trichopoda* as out-group and the concatenated translated nucleotide alignments of 230 single-copy orthologues were used as input of MCMCtree. The phylogeny was calibrated using various fossil records or molecular divergence estimated by placing soft bounds at split node of (MYA): *A. trichopoda* and *O. sativa* (’>1.73<1.99′), *D. carota* and *Salvia splendens* (’>0.95<1.06′), *Arabidopsis thaliana* and *Populus trichocarpa* (’>0.98<1.17′) obtained from the TimeTree database (Kumar et al., 2017).

### Synteny, Positive selection genes and WGD anaylsis

The genomic alignment between *C. burmannii* and *C. micranthum* was finished and visualization by MUMMER (Kurtz et al., 2004). We performed multiple sequence alignment to extract the conserved paralogs of the protein sequences of this data by using BLASTP (E-value ≤ 1e−5). MCScanX was used to identify collinearity blocks, and Ks values were calculated by software WGDI to identify and extract collinear interval and corresponding gene pairs (Sun et al., 2021). The calculation of Ks with the method YN00 was implemented to predict the Whole Genome Duplication (WGD) events

### Transcriptome profiling analysis

Pipeline Hisat2 was reads alignment of the reads from RNA-seq and the read counts matrix was generated by software featureCounts (Liao et al., 2014) for the quantitative analysis expression. R package DESeq2 was implemented to normalize the read counts data and identified the DEGs between different organs with Pvalue ≤ 0.01 and an absolute log2 (fold change) ≥1 as the threshold (Love et al., 2014). Software STEM was utilized to the cluster analysis DEGs expression and GO enrichment analysis (Ernst and Bar-Joseph, 2006).

### Tps gene family anaylsis

Proteome data sets including *C. burmannii* and other four sets *C. micranthum, M. acuminate, O. sativa* and *D. carota* from NCBI were utilized in TPS gene family analysis. Pfam domains: PF03936 and PF01397, were used to identify against the proteomes using HMMER (version 3.0; cut-off at e-10) (Finn et al., 2010; Finn et al., 2011). Putative or annotated protein sequences of *TPS* were aligned using MAFFT with default parameters and manually adjusted using MEGA. The TPS gene tree was constructed using iqtree with 1,000 bootstrap replicates. The subfamily *TPS*-a was designated as the out-group. Branching nodes with bootstrap values of <80% were treated as collapsed.

### Prediction of protein structure modeling and molecular docking

Homology modeling was performed with Rosettafold with default *de novo* prediction parameter (Baek et al., 2021). The results were inspected and rendered with PyMOL (DeLano, 2002). Protein docking and binding energy calculation were done with AutoDock Vina using local search parameters and default docking parameters (Trott and Olson, 2010). The alignment and the visualization of amino acid sequences was finished by platform ClustalW and ESPript (Gouet et al., 1999).

### Data and code availability

The sequencing data, including the assembly data of *C. burmannii* was submitted to the National Genomics Data Center (NGDC) under BioProject accession number PRJCA006286. All the assembly and annotation data also can be accessed from CAMD (http://www.cinnamomumdatabase.com/)

Customized codes used in this study have been deposited in GitHub https://github.com/lifangpings/CB_genome.

### Real-time Quantitative PCR (q-PCR) comfirmation

For q-PCR assays, the total RNA was extract using Aurum Total RNA Mini Kit (BIO-RAD) and cDNA was obtained by thermo scientific RevertAid First Strand cDNA Synthesis Kit according to the supplier’s protocols respectively. All the primers’ sequence was obtained from the National Centre for Biotechnology Information (NCBI) database. According to kit instruction 10 μl reaction system was prepared and q-PCR was operated using CFX96 Real-Time System (BIO-RAD). For data analysis, all detected genes expression were normalized against Actin-F (CBB03T203270) of *C. burmannii*.

### Database construction

All genomic sequence, annotation, were stored via MySQL on a Centos7 server. A user-friendly website was developed using HTML5, JavaScript and PHP7, which can be accessed through different browsers, such as Google Chrome and Firefox. Gene models and transcript isoforms were provided via JBrowse (Skinner et al., 2009; Buels, R. et al., 2016). The query searches were achieved via PHP7. Common utilities for genomic studies such as BLAST, and Sequence exactor were also deployed and accessible.

## Result

### Genomic assembly

To evaluate the genome size and heterozygosity of *Cinnamomum burmannii* (chvar. Borneol), a total of 61.4 GB short reads from the MGISEQ-2000 sequencing platform were subjected to K-mer analysis. The 17-mer frequency curve showed a unimodal distribution, with the highest peak occurring at a depth of 54 (Supplementary Fig. S1a). Based on the total number of K-mers, the genome size and heterozygosity of *C. burmannii* (chvar. Borneol), were estimated to be 0.94-1.25 GB and ~0.67%, respectively (Supplementary Fig. S1b; Supplementary Table. 1). A total of 170.1 GB single molecule sequencing long read data with the average length about 32 kb (N50=21.38kb) was utilized in the genomic *de novo* assembly. After polishing combining the short-read data and removing redundant/contaminated sequences, the size of the final contig was 1.15 GB with a contig N50 of 1.13Mb (Table.1). A Hi-C interaction heatmap indicated 12 highly obvious interacted range of the signal which is corresponding with the previous research about the chromosome number of *C. burmannii* (chvar. Borneol) (Supplementary Fig. S2). Based on the signal, a total of 98.77% of the assembly was anchored to 12 pseudo-chromosomes. The contig sequences were connected in the determined order and direction by adding 100 N to obtain the final chromosome-level genome sequence with a chromosome mount rate of 98.77% the size ranged from 66.1Mb to 140.5Mb, and with Scaffold N50 = 94.90Mb. The ordering, and orientation of the contigs was also arranged, providing the first high-quality chromosome-scale genome assembly for *C. burmannii* (Table.1).

**Table.1.**
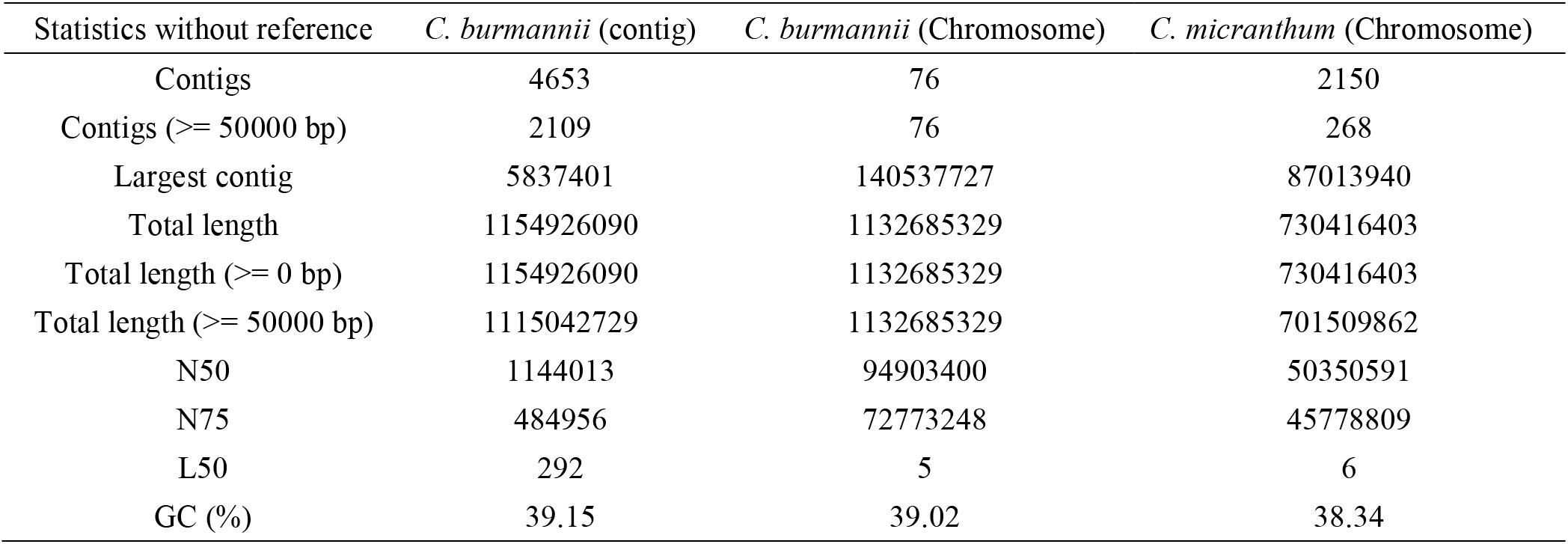
Summary and comparison of *C. burmannii* and *C. micranthum* assembly

| Statistics without reference | *C. burmannii* (contig) | *C. burmannii* (Chromosome) | *C. micranthum* (Chromosome) |
| --- | --- | --- | --- |
| Contigs | 4653 | 76 | 2150 |
| Contigs ($\geq 50000$ bp) | 2109 | 76 | 268 |
| Largest contig | 5837401 | 140537727 | 87013940 |
| Total length | 1154926090 | 1132685329 | 730416403 |
| Total length ($\geq 0$ bp) | 1154926090 | 1132685329 | 730416403 |
| Total length ($\geq 50000$ bp) | 1115042729 | 1132685329 | 701509862 |
| N50 | 1144013 | 94903400 | 50350591 |
| N75 | 484956 | 72773248 | 45778809 |
| L50 | 292 | 5 | 6 |
| GC (%) | 39.15 | 39.02 | 38.34 |

Multiple method was utilized for the quality assessment of the novel *C. burmannii* genome. The method of Benchmarking Universal Single-Copy Orthologs (BUSCOs) was implemented to evaluate the assembly quality of *C. burmannii* genomes, which identified 89.7% of the complete BUSCOs in the assembly by using the reference of the conservative database of dicotyledons (Supplementary Table S2). For the assessment of genomic consistency, the short reads were mapped to the genome using bwa software. This analysis indicated ~99.78% of the clean data were mapped to the genome assembly. The mapping ratio of the RNA-seq reads from different tissues was in the range of ~92.08%–94.93%. The GC content of this genome is 39.02% while the sequencing depth was concentrated at 110–150×. All these methods of assessment elucidated a high completeness and consistency of the *C. burmannii* genomic assembly (Table.1).

### Repeat element identification and Genome annotation

The *de novo* prediction based on RepeatModeler indicated 673.32 Mb of repetitive sequences, accounting for 63.24% of the assembled genome. Combining the result of *de novo*, homolog-based, and transcriptome-based predictions, a total of 41549 non-redundant protein-coding genes were generated across 12 chromosomes (Supplementary Fig. S3). The average gene length and coding sequence size were 2962.35 and 1142.14 bp, respectively, with an average of five exons per gene. The result of function annotation indicated that 37799 genes (90.97 %) was annotated in at least one pathway of the public database. Further functional annotation using InterProScan estimated that of the genes 33582 (80.82%) contained conserved protein domains recorded by Panther and Pfam database. 54.38% of the genes could be classified with Gene Ontology (GO) terms. 8401 (20.22%) of the genes mapped to known plant biological pathways based on the KEGG Pathway database.

### Phylogenetic analyses

To identify the phylogenetic relationship and position of *C. burmannii* and *C. micranthum* we constructed a phylogenetic tree based on 230 strictly single-copy orthologous sets. Consistent with expectations, phylogenetic tree indicated clustering of two species of *Cinnamomum* on one branch. The maximum likelihood (with a high bootstrap index) and MCMCtree with fossil calibrations analysis indicated that the divergence between magnoliids and eudicots tended to happen in 201 Mya (95% confidence interval: 196.1-209.1). The time of divergence between *C. micranthum* and *C. burmannii* to be 8 Mya (95% confidence interval: 7.5-8.5) (Figure. 1).

**Figure.1.**
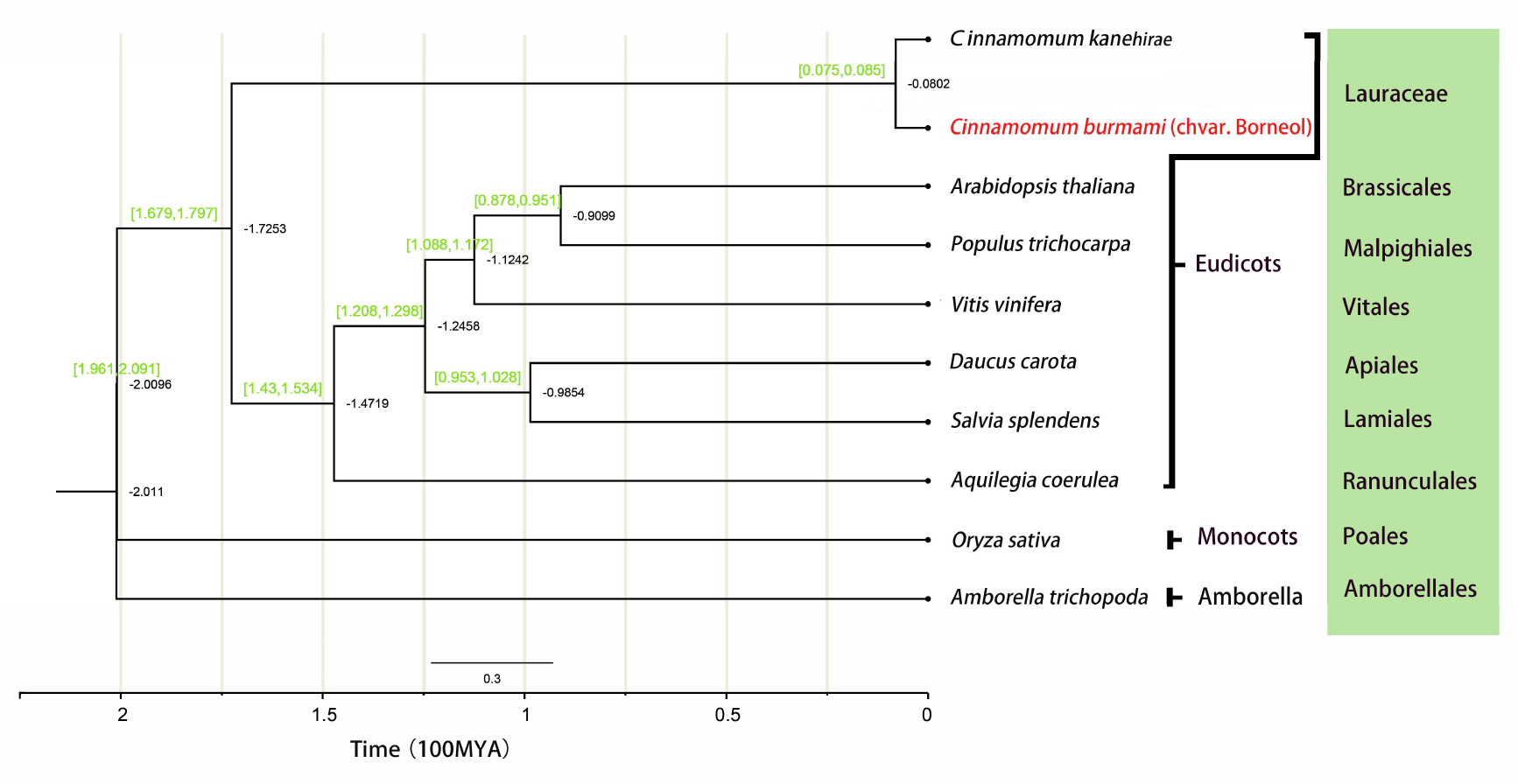
The phylogenetic relationship of *C. burmannii, C. micranthum* and other eight plant species of different family. The green numbers in the brackets denote divergence time estimates. All nodes’ bootstrap support was 100 unless stated otherwise.

### LTR-RTs promote species differentiation among Cinnamomum

In the identification of LTR-RTs for these two species, a total of 186.62Mb (25.1% of the whole genome) LTR-RTs sequence was detected in *C. micranthum* while 459.42Mb (40.59% of the whole genome) was detected in *C. burmannii* (Figure. 2a and Supplementary Table S2). The insert time analysis elucidated these two species experienced a significant recent outbreak of LTR-RTs insertions. However, much more insertions of LTR-RTs were speculated to occur in 0-8Mya in *C. burmannii* than in *C. micranthum,* which was also corresponding with the divergent time between these two species. These LTR-RTs were also evenly distributed in the genome of *C. burmannii* (Figure. 2b).

**Figure. 2.**
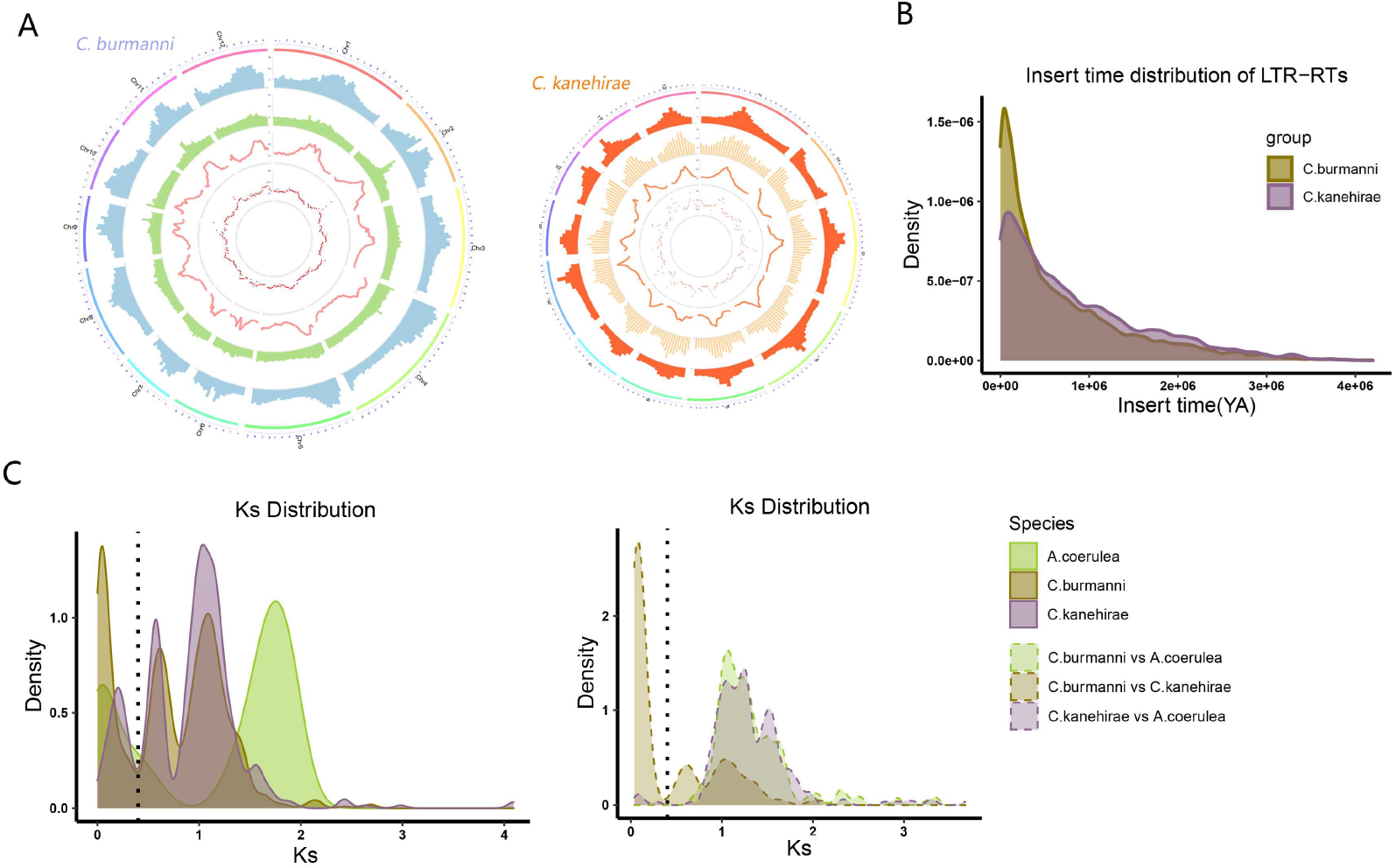
The distribution and insertion times of the LTR-RTs in *C. burmannii* and *C. micranthum* genomes as well as the WGD event identification. **A:** LTR-RTs Characterization of the *C. burmannii* and *C. micranthum* genome; the circle from outside to inside represents chromosomes LTR-Copia density, LTR-Gypsy density, Total LTR density and GC content All distributions are drawn in a window size of 3 Mb. **B:** Quantity and time distribution of LTR-RTs **C:** Ks distribution among *C. burmannii*, *C. micranthum* and *A. coerulea* Lines show Ks distribution within (continuous) and between genomes (dashed).

This ~250 Mb length difference of LTR-RTs tends to be a preliminary explanation for the interspecific genome size differences in *C. micranthum* and *C. burmannii* genera.

### Whole-genome duplication and Synteny analysis

To more precisely infer the timing of the two rounds of WGD evident in the *Cinnamomum,* the intragenomic and interspecies homologue *Ks* (synonymous substitutions per synonymous site) distributions were estimated with the genome of *C. micranthum, C. burmannii* and *A. coerulea* (as outgroup). The curve of *Ks* distribution indicated that two peaks at ~0.5 to 1.5 in both *C. micranthum* and *C. burmannii* genomes, which elucidated these two species shared the same two ancient WGD events (Two ancestral WGD events endemic to Lauraceae before divergence of eudicots and magnoliids) before their divergence and no recent independent WGD event was detected (Figure. 2c).

Consistent with the predictability, both the results of whole genome comparison and gene collinear comparison between *C. micranthum* and *C. burmannii* indicated an obvious synteny presence (Figure. 3a and Supplementary Fig. S4). Based on the collinearity of *C. burmannii* and *C. micranthum* genomes, we sorted and renamed *C. burmannii* genome sequences. We found that the relative proportions of chromosome sizes were consistent between the two species (Table. 2). Besides, large fragments of chromosomal translocations were displayed in Chr2, Chr 4, Chr5, and Chr12, while a large fragment inversion occurs in front of Chr6 in comparison between this two *Cinnamomum* species. This phenomenon suggested that the translocations and inversion of chromosomes might have been occurred during the process of *Cinnamomum* species differentiation.

**Figure. 3.**
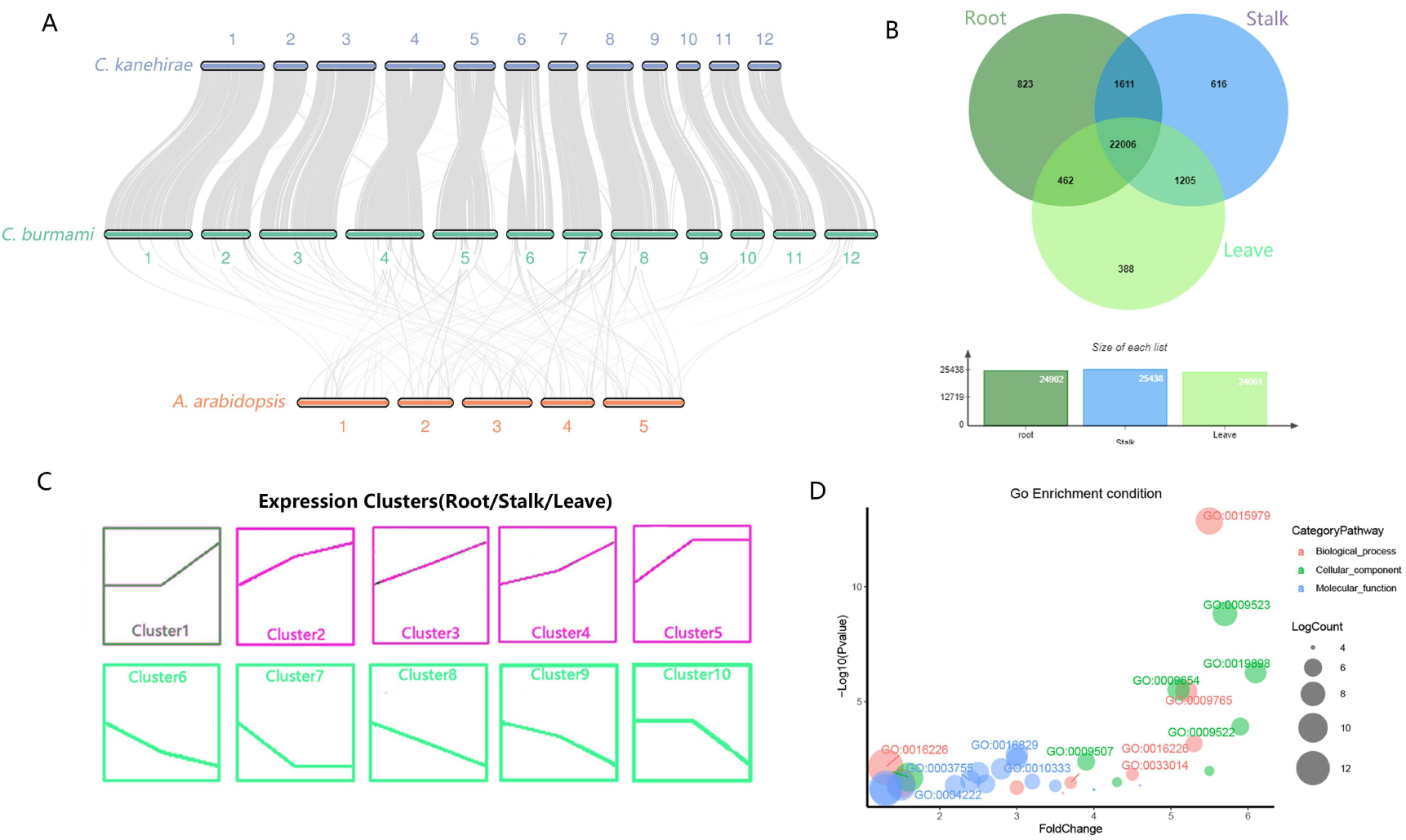
Collinearity analysis between *C. burmannii, C. micranthum* and *A. Arabidopsis* as well as the gene’s organ-specific expression of different organs *of C. burmannii.* **A:** Gene Collinear relationship of *C. burmannii* and *C. micranthum* and *A. arabidopsis*. The gray line connects matched gene pairs. **B:** The Venn diagram visualized the expressed genes in roots, stalks, and leaves. The overlapping regions represent genes expressed in at least two tissues, while the non-overlapping regions represent tissue-specific genes. **C:** Hierarchical clustering showing the expression patterns of DEGs in roots, stalks, and leaves.

**Table.2.** Statistics of the sizes of each chromosome of *C. burmannii* and *C. micranthum*

| Chromosome | *C. burmannii* | *C. micranthum* | Corresponding Chromosome No. (*C. micranthum*) |
| --- | --- | --- | --- |
| Chr1 | 140537727 | 87013940 | QPKB01000001.1 |
| Chr2 | 92646709 | 50350591 | QPKB01000006.1 |
| Chr3 | 122226000 | 81065106 | QPKB01000002.1 |
| Chr4 | 135862046 | 77926444 | QPKB01000003.1 |
| Chr5 | 108037362 | 58596509 | QPKB01000005.1 |
| Chr6 | 68311217 | 48047431 | QPKB01000007.1 |
| Chr7 | 56091752 | 39555212 | QPKB01000011.1 |
| Chr8 | 94903400 | 58831143 | QPKB01000004.1 |
| Chr9 | 66575986 | 41984183 | QPKB01000010.1 |
| Chr10 | 66169134 | 37069385 | QPKB01000012.1 |
| Chr11 | 72773248 | 45778809 | QPKB01000009.1 |
| Chr12 | 89292919 | 46631104 | QPKB01000008.1 |

### Characteristic analysis of genes showing organ-specific expression

A total of 31537 genes were detected to expression in the organs of *C. burmannii*. The gene expression levels analysis indicated that 1318, 1208, 672 genes expressed explicitly in roots, stalks, leaves respectively, while 24681 genes were expressed in all tissues (Figure. 3b).

To elucidate the similarities and differences of gene expression patterns in different tissues, we performed a k-means cluster analysis. A total of 20444 differentially expressed genes (DEGs) were divided into 10 clusters. We focused on clusters containing tissue-specific expressed genes, particularly those that were stable in roots and stalks while increased in leaves (3348 genes in Cluster1) (Figure. 3c). Apart from the genes involving in photosynthesis, genes in the terpene synthase (*TPS*) activity pathway (GO: 0010333) were shown to be expressed and enriched in leaves, which was consistent with the higher content of terpenoids in leaves including D-borneol in *C. burmannii* reported by previous studies. Besides, a number of methylation related genes (GO: 0004222 and GO: 0008173) were also significantly increased and enriched in leaves. This phenomenon suggested the regulation associated with epigenetic may play a role in development and compound synthesis in the leaves of *C. burmannii* (Figure. 3d).

### TPS gene family analysis

Similar to another plant of *Cinnamomum, C. micranthum*, a large number of *TPS* genes (*CbTPS*) located in the genome of *C. burmannii.* A total of 73 *CbTPS* genes were predicted and annotated. Phylogenetic analysis of *CbTPS* placed these genes into six *TPS* gene subfamilies that have been described for seed plants. Among this *CbTPS* genes, 1 and 2 were placed in the *TPS-c* and *TPS-e* subfamily respectively. These two subfamilies have been reported to encode diterpene synthases like copalyl diphosphate synthase and *ent*-kaurene synthase (Chaw et al., 2019). These are key enzymes catalyzing the formation of the 20-carbon isoprenoids (collectively termed diterpenoids; C20s). The remain *CbTPS* genes attributed to *TPS*-a, *TPS*-b, *TPS*-f and *TPS*-g, which probably encode the 10-carbon monoterpene (C10) synthases, 15-carbon. With 29 and 23 homologues, respectively, *TPS*-a and *TPS*-b subfamilies are most diverse in *C. burmannii*, presumably contributing to the mass and mixed production of volatile C15s and C10s, including D-borneol (Figure. 4 and Supplementary Table S4).

**Figure. 4.**
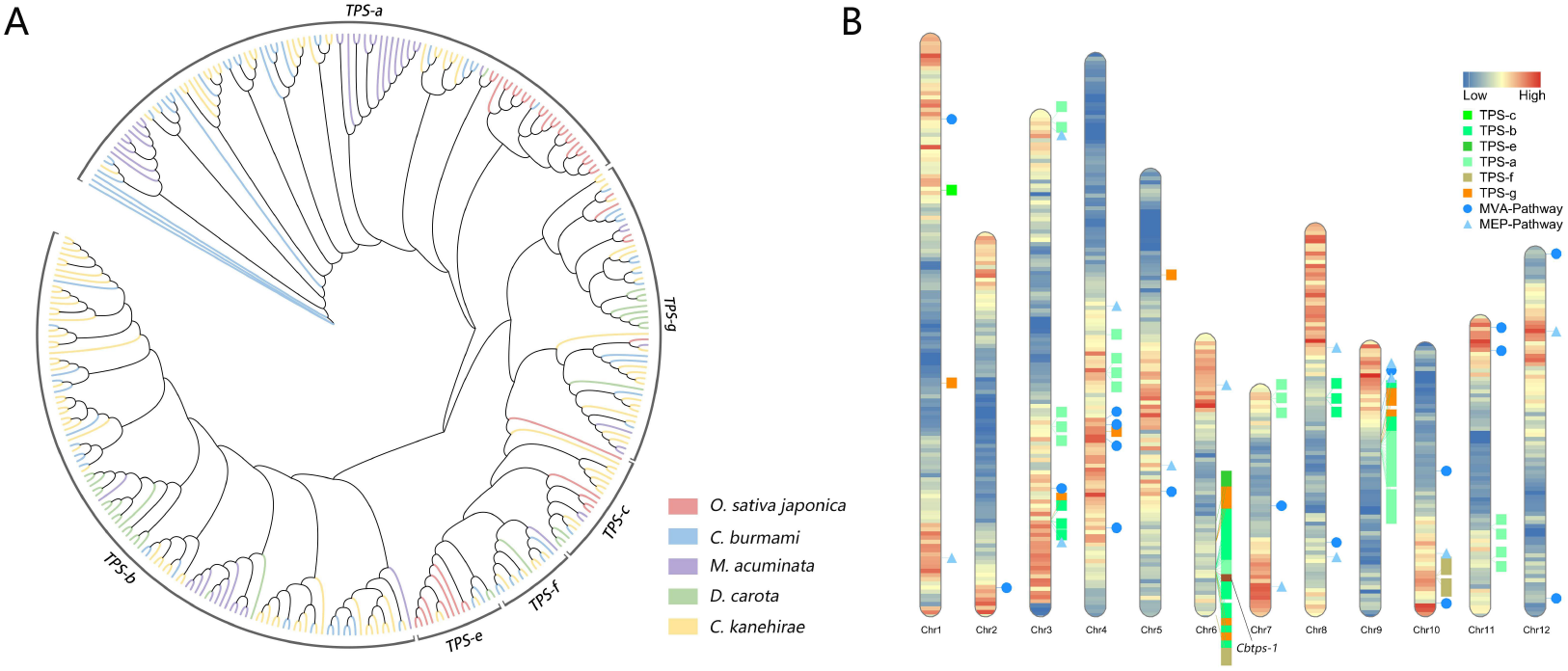
The gene density and the gene distribution involving in *TPS* gene family and D-borneol formation; Phylogenetic placements of the 73 *TPS* family genes of *C. burmannii***. A:** The phylogenetic tree was constructed using putative or characterized *TPS* genes from 5 plant genomes including *C. burmannii*. **B:** Gene density distribution as well as the location of the gene involving in *TPS* gene family and D-borneol formation.

Besides, *CbTPS* genes are not uniformly distributed throughout the chromosomes and clustering of members from individual subfamilies was observed as tandem duplicates. Among these genes, forty three copies (58.9%) of *CbTPS* genes belonging to different subfamilies were found in the 53–59Mb region of chromosome 6, and 56–61Mb region of chromosome 9. Two to nine *TPS* family genes were detected in tandem on chromosomes 1, 3, 4, 7, 10 and 11 while only one gene of *TPS* family was detected in chromosomes 5. chromosomes 7 contains 29 *CkTPS* genes belonging to several subfamilies, including all of the eight *CkTPS*-a, 12 *CkTPS*-b, five *CkTPS*-e and three *CkTPS*-f Genes belonging to this cluster were not grouped together in their corresponding subfamily phylogen, suggesting that their arrangement might have occurred more recently than the last WGD event (Supplementary Fig. 4b).

### Identification of genes related to the D-borneol biosynthesis pathway

Previous research indicated that MVA pathway play a key role in the upstream biosynthesis pathway of D-borneol. Our gene annotation results indicated that all MVA pathway genes (*CbERG10, CbERG13, CbtHMG1, CbERG12, CbERG8, CbERG19, CbIDI1, CbERG20*) distributed in 12 chromosome of *C. burmannii.* The transcriptome analysis elucidated the expression level of genes *CbERG20, CbIDI-1,* was significantly elevated in leave comparing to stem and root. *CbERG20* is the key enzyme to promote the synthesis of GPP, which is a directly related raw material of D-borneol synthesis.

Besides, MEP pathway also have been reported to participate in the precursor GPP formation in the D-borneol formation process. In this pathway, the expression of multiple genes such as *CbCMK, CbMCS* and *CbHDR* showed similar expression pattern with above key genes in MVA pathway. (Figure. 5).

**Figure. 5.**
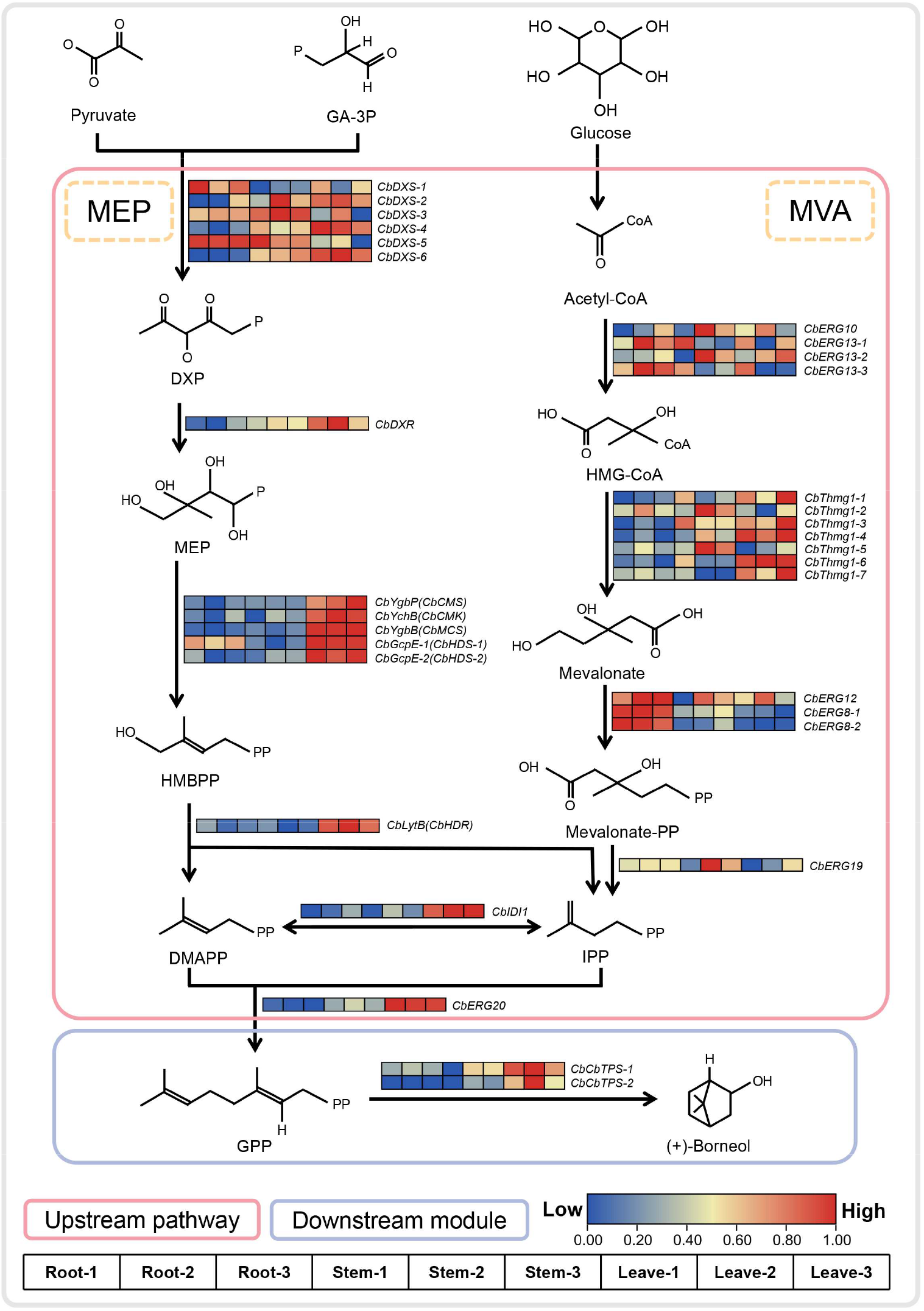
Identification and expression analysis of genes involved in MVA and MEP pathway for key material GPP production (pink ellipses) nd the biosynthetic pathway of D-borneol (blue ellipses). Heat maps show the expression of genes involved in these pathways

For the downstream pathway, *CbTPS-1* (*CBB06T352390*), encoded by a gene of *TPS*-b gene subfamily, has been reported to directly mediate the transformation of GPP to D-borneol (Ma et al., 2021). The expression levels of these key genes for D-borneol synthesis were much higher in leaves than in stem or root implied they may be the key or the limiting factor in the biosynthesis of D-borneol in *C. Burmannii* leaves (Figure. 5). Gene cloning and qPCR analysis to above potential key genes were further conducted, and results are highly consistent with our transcriptomic and gene annotation results, indicating the accuracy and reliability of our results (Supplementary Fig. S6).

### Sequences differences cause configuration alteration of the docking site of CbTPS-1 and CkTPS-1

The 3D structure model of protein generated by Rosettafold was implemented to model the protein structure of *CbTPS-1* and its highly homologous *CkTPS-1*(*RWR98053.1*) in *C. micranthum*, and molecular docking analysis with small micromolecule GPP was also performed. This analysis showed an obvious GPP binding pocket in the middle of these two *TPS* proteins folding structure (Figure. 6a). The GPP docking pocket located in the ~300aa to ~500aa of these two TPS protein. A total of 26 amino acid mutations were detected in this region. Among these, 15 mutations changed the polarity of the amino acids at that location, which have potential to affect their protein folding (Figure. 6b). Therefore, due to folding differences caused by changes in amino acid sequence, GPP can form more hydrogen bonds with more A-helices (4 > 1) in *CbTPS-1* binding pockets (9 > 8) and its binding energy in *CbTPS-1* is also higher than that of *CkTPS-1* (−7.7<−7.4). The higher binding capability of *CbTPS-1* may cause of higher D-borneol production in *C. Burmannii* than other These differences may be the main cause of *C. Burmannii* to produce higher D-borneol than other *Cinnamomum.* plants (Figure. 6c).

**Figure. 6.**
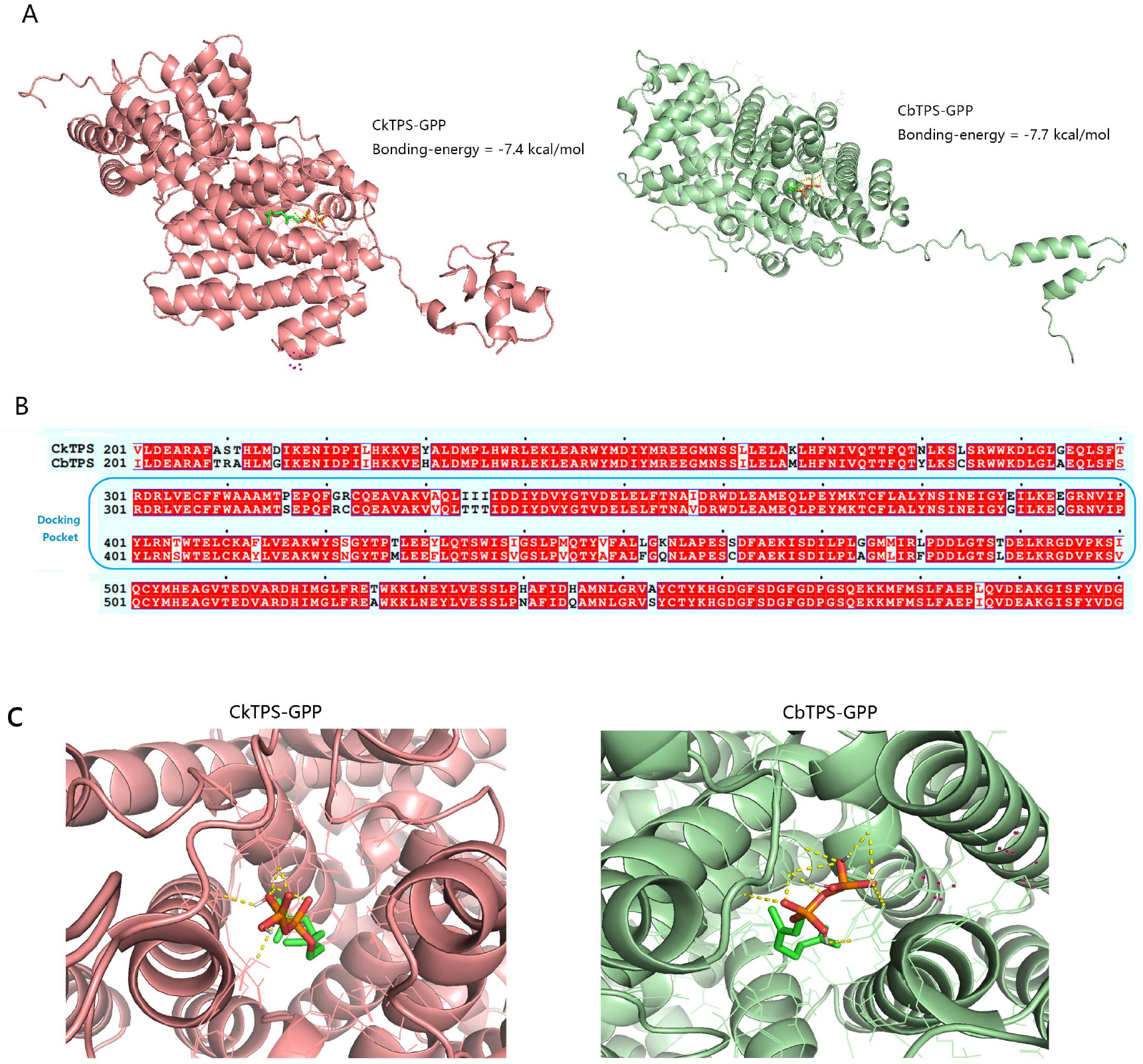
Molecular docking result between *CbTPS-1* and its highly homologous gene *CkTPS-1* in *C. micranthum.* **A:** 3D structure of proteins and the overall docking situation (docking pocket position; *CkTPS-1*-GPP/*CbTPS*-1-GPP); **B:** The sequences comparison of Amino acid docking pocket position **C:** The details of docking (*CkTPS-1*-GPP/*CbTPS-1*-GPP).

### Construction of genome database of Cinnamomum species

To meet demand for pitaya genome and multiomics data resources, we established an integrated pitaya genome and multiomics database (CAMD; http://www.CinnamomumDatabase.com). The CAMD provides a comprehensive consultation service including genomic blast, gene browser and sequence fetch with the data from the latest assemblies of *C. burmannii* and *C. micranthum* genomes (Chaw et al., 2019) (Figure. 7).

**Figure. 7.**
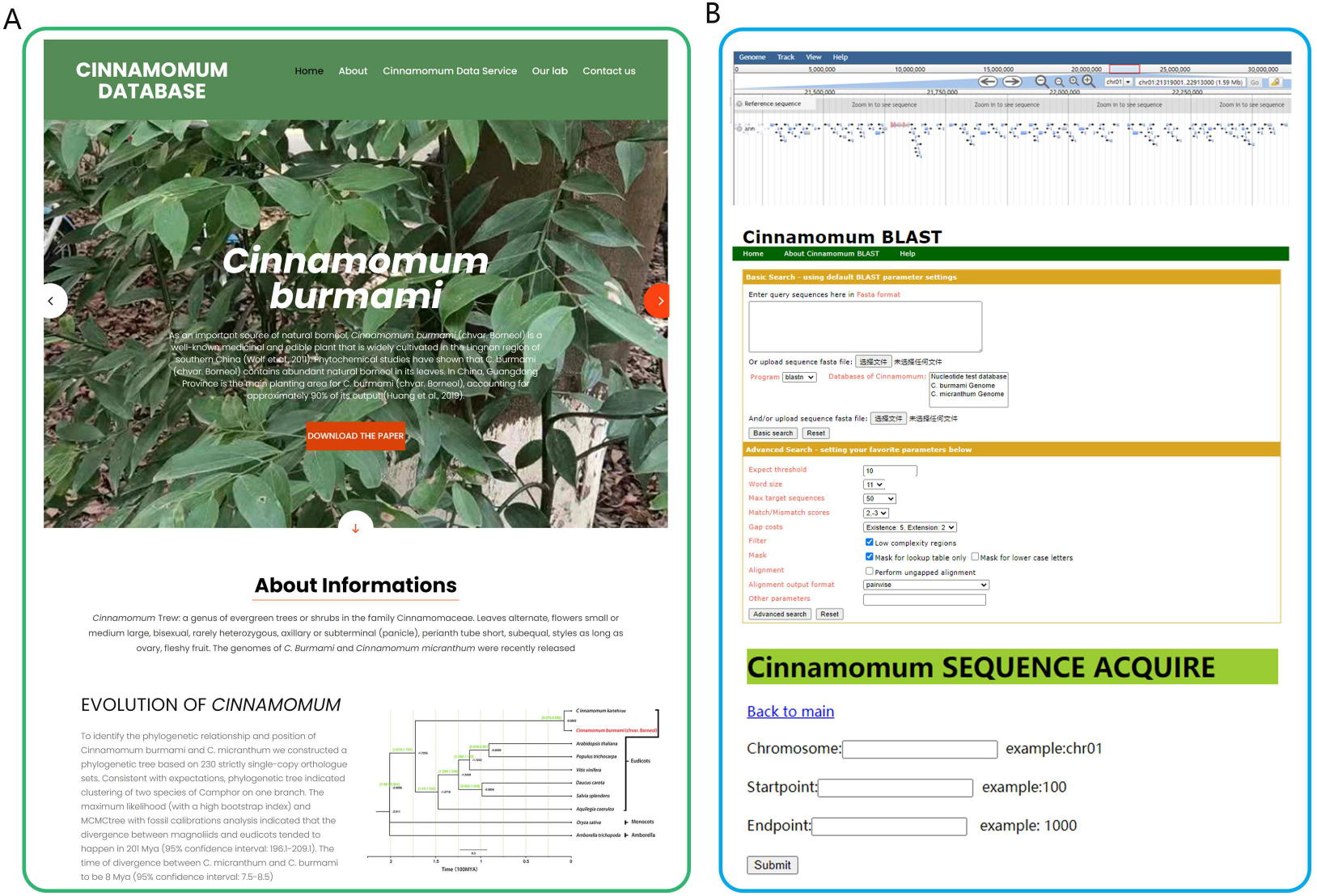
The screenshots of representative resources and action page example of CAMD; **A:** Main pages of CAMD; **B:** Partial function pages displayed including Genomic blast, Gene Brower and sequence fetch.

## Discussion

*Cinnamomum*, a genus of about 250 species of the Lauraceae, is widely distributed in tropical Asia, Australia to Pacific islands, and tropical America. The plants of *Cinnamomum* are rich in volatile oil, terpenoids, tannins and their analogues, aromatic compounds and polysaccharides. Multiple species of this genus are important raw materials in chemical industry and medicine. However, the only one assembled genome in this species is *C. micranthum* (Chaw et al., 2019). There are few reports on further genome research of *Cinnamomum* genus. The lack of high-quality genome and detailed genomic information has limited our understanding of molecular mechanisms underlying valuable natural product and genetic improvement of these plants.

In the present study, we assembled a chromosomal scale, high-quality genome of *Cinnamomum burmannii* (chvar. Borneol), as well as detailed genomic and transcriptomic information. Compared with the first published genome of *Cinnamomum species, C. micranthum,* the assembly of *C. burmannii* achieved much higher quality, such as a longer sequence length of contigs (1.14 Mb > 498.92Kb) and a higher chromosome mount rate (98.31% > 90.1%) with a larger genomic size (1.13Gb > 730.7 Mb). These improvements suggested that the assembly of *C. burmannii* tends to be a valuable and better reference genome for future genomic study of *Cinnamomum* species.

The construction of the *C. burmannii* genome and the comparative genome analysis also shed light on the understanding of genome evolution of Lauraceae. Our results confirmed that Lauraceae and camphor plants shared two ancestral WGD events. As the representative of magnoliid, the phylogenetic relationship analyses of *C. burmannii* also resolve magnoliids to be closer to eudicots than to monocots. Furthermore, the outbreaks of LTR-RTs have been considered to be an important impetus of increasing genetic diversity of multiple species (Sun et al., 2020; Lin et al., 2018). The combination of LTR-RTs size and insertion time indicated that recent LTR-RTs insertion events also promoted the differentiation between species *C. micranthum* and *C. burmannii* in *Cinnamomum.*

Our assembled genome and transcriptomic analysis results also help to elucidate the molecular mechanism of borneol biosynthesis. As a medicinal plant, *C. burmannii* (Chvar. borneol) is an important source of D-borneol. D-borneol, which is also called natural borneol, is one of the most famous Chinese traditional medicines. It has the effect of awakening, clearing heat and relieving pain. It is often utilized popularly for fever, dizziness, convulsion, stroke, phlegm, sore throat, purulent ear canal and other similar symptoms. Since its discovery in the 1980s, *C. Burmannii* (Chvar. borneol), which is rich in natural borneol, has been widely grown as a medicinal crop in city Meizhou in Guangdong province of China. It is very important to fully elucidate the molecular pathway of D-borneol biosynthesis for further natural plant utilization and future synthetic biology application. Ma et.al had reported the reconstitution of D-borneol biosynthetic pathway in *Saccharomyces cerevisiae*, which is a critical step for borneol biosynthesis application. However, genes responsible of borneol synthesis, and theirs underlying regulation mechanisms in plants are still largely unknown.

With the help of high-quality reference genome of *C. burmannii* (Chvar. borneol) assembled in the present study, we were able to fully dissect the natural borneol synthesis pathway from a genome scope. Previous studies have confirmed MVA pathway is the key pathway for the synthesis of GPP, which serve as the precursor of D-borneol (Ma et al., 2021). The MEP pathway has also been reported to be involved in GPP synthesis in plants (Banerjee and Sharkey, 2014). The transformation of GPP and D-borneol was completed with the participation of *TPS* family, while *CbTPS-1* has been confirmed to participate in D-borneol biosynthesis directly (Ma et al., 2021). In the present study, all genes participate in the both natural borneol biosynthesis pathway were identified in *C. burmannii* (Chvar. borneol) genome. Transcriptomic analysis showed key genes in both pathways tends to highly expressed in leaves, rather than in stem or root. All these results provided important genomic information of genes for the following research about D-borneol biosynthesis in *C. burmannii*.

Furthermore, the analysis of *TPS* gene family elucidated multiple *TPS* gene clusters were detected in *C. burmannii* genome, especially in Chr6 and Chr9. Recent studies have shown that several plants biosynthetic gene clusters (BGCs) of natural products exist in plant genomes. These plant biosynthetic gene clusters generally participate in the process of natural product synthesis synergistically (Liu et al., 2020; Mao et al., 2020). In the present study, the function and mechanisms of BGCs formed by *TPS* family genes are worth more profound exploration.

Additionally, our transcriptomic analysis further identified genes participate in methylation process were significantly enriched in genes specific expressed in leaves, but not in stem or root. Several researches have reported the epigenetic processes, such as DNA or RNA methylation played important roles in the synthesis of secondary metabolites in multiples species (Zuo et al., 2020; Wang et al., 2019). Leaves is the major organ for borneol synthesis in *C. burmannii*. Together with the results that expressions of genes of borneol biosynthesis pathway were much higher in leaves than in stem and root, and the results that methylation related genes enrichment in leaves specific expression genes, implied methylation may play important roles in the expression regulation of gene related to borneol synthesis in *C. burmannii* leaves. Methylation and gene expression regulation may be further leverage as target for borneol production improvement in *C. burmannii* (Chvar. borneol). The mechanisms behind these results need further investigation.

In the present study, we also constructed a genome database (CAMD), which combined our results of *C. burmannii.* and genome information from another *Cinnamomum* plant, *C. micranthum.* (Chaw et al., 2019). CAMD is the first database of the *Cinnamomum* and will become an excellent central gateway to better understand the biology and genetics of Lauraceae. The release of CAMD will benefit for studies of genetic diversity and improvement in *Cinnamomum*.

## Conclusion

Here we presented a high-quality complete genome, as well as detailed genomic information of *C. burmannii* (Chvar. borneol). The high-quality genome provides novel insights about genome evolution of *C. burmannii*, as well as *Cinnamomum* genus. With the help of assembled and annotated genome, we dissected the pathways of borneol synthesis, and identified key genes in these pathways. Transcriptomic analysis further shed light on the further investigation of regulation and molecular mechanisms of these pathways. We also constructed the genome database (CAMD) of *Cinnamomum* species for better data utilization in the future. The availability of *C. burmannii* complete genome and its genomic information from this work will provide a valuable genome resource for understanding the evolution, active ingredient biosynthesis, and genetic improvement for not only *C. burmannii* itself, but also *Cinnamomum* genus.

## Acknowledgements

This work was supported by the scientific innovation strategy-construction of high-level Academy of Agriculture Science (R2019PY-JX003 and R2019PY-JX001), Natural Science Foundation of Crops Research Institute, Guangdong Academy of Agricultural Sciences (0145). We thank Prof. Wang Shaokui (Department of Crop genetics and breeding, South China Agricultural University) and Prof. Zhou Xiaofan (Integrative Microbiology Research Centre, South China Agricultural University) for kindly help and support.

## Author contributions

LF, ZJ and WJ conceived and designed this research. LF, HS, XS, WB, HZ, ZP, MY, ZJ, LQ, HZ and WJ performed the experiments. LF, HS, WB, ZJ and WJ prepared the article.

## List of Supplementary materials

### Supplementary Figure

Supplementary Figure.1 Evaluation of genome size and heterozygosity

Supplementary Figure.2 HI-C heatmap of *C. burmannii*

Supplementary Figure.4 Collinearity of genomic sequences between *C. burmannii* and *C. micranthum*

Supplementary Figure.5 qPCR confirmation of partial gene involving in D-borneol biosynthesis in *C. burmannii*

Supplementary Figure.6 Gene clone of *CbERG20* and *CbTPS-1* from *C. burmannii*

### Supplementary Table

Supplementary Table.1 Genomic survey result of *C. burmannii* generated by software Gce

Supplementary Table.2 Busco assessment results of *C. micranthum*

Supplementary Table.3 Length and types Statistics of LTR-RTs

Supplementary Table.4 Numbers of *TPS* subfamilies in the 5 genomes

Supplementary Table.5 List about the gene in *TPS* family or involving in D-borneol biosynthesis in *C. burmannii*

Supplementary Table.6 Sequence and List of primers in qPCR and gene clone confirmation assays.

