## Supplementary material for "High-quality genome assembly of *Cinnamomum burmannii* (chvar. Borneol) provides insights into the natural borneol biosynthesis": sp

#### Supplementary Figure:

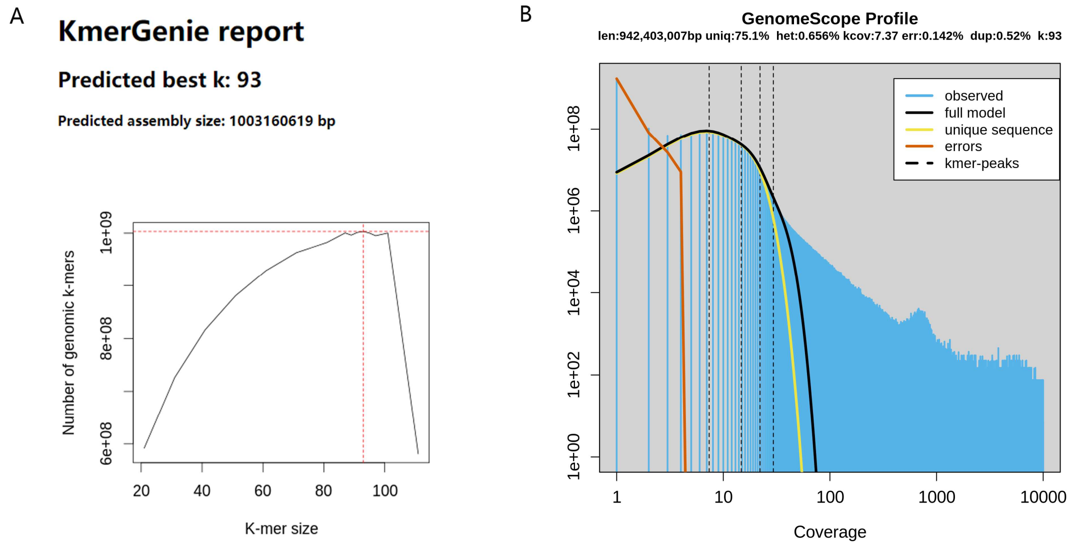


#### Supplementary Figure.1

**Evaluation of genome size and heterozygosity A:** Genomic size and suitable K-mer assessment of *C. burmanni* genome; **B:** Evaluation of *C. burmanni* genome size and heterozygosity

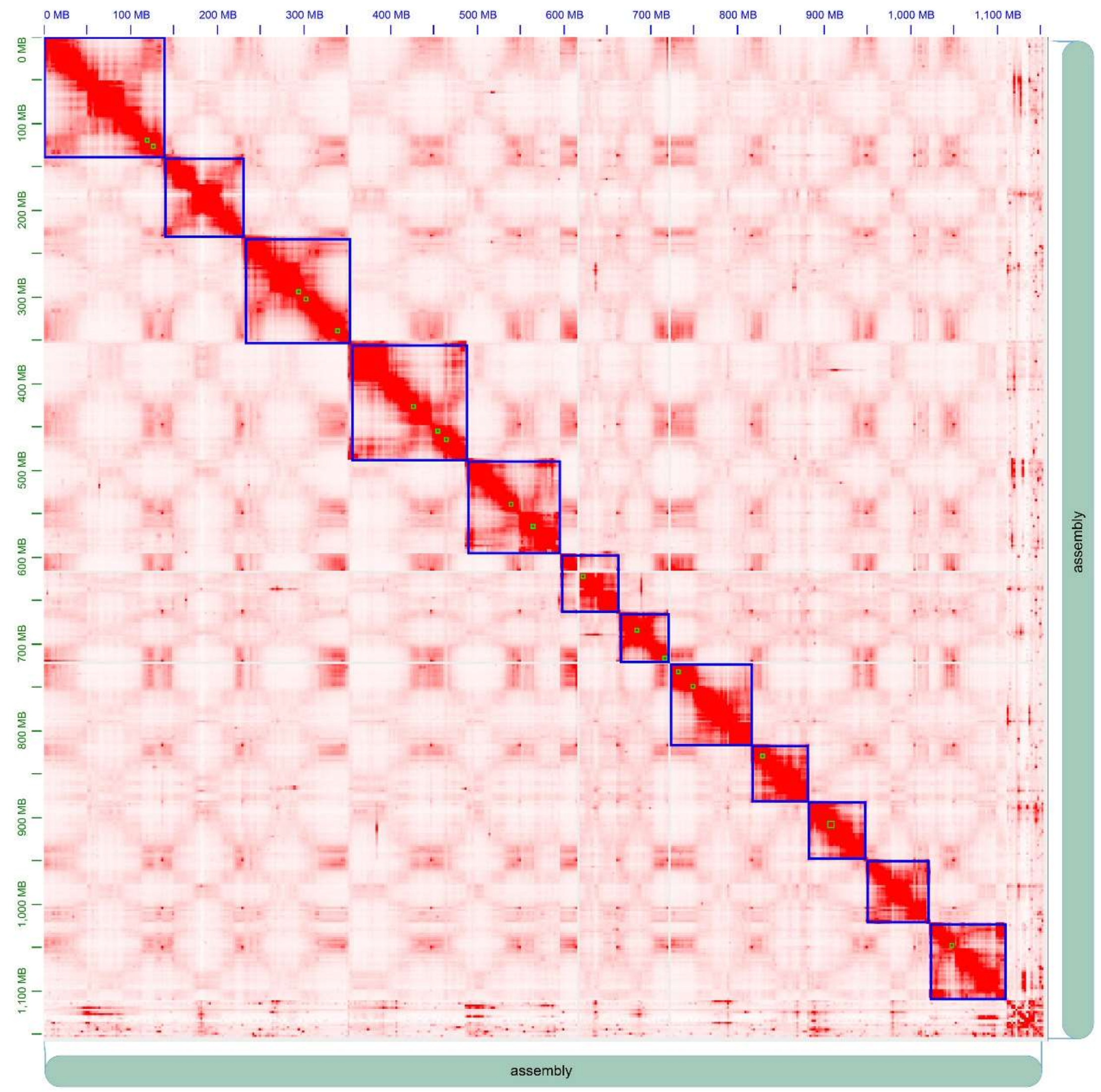


**Supplementary Figure.2**

**HI-C heatmap of *C. burmanni;*** 12 Twelve distinct areas of interaction displayed in this heatmap.

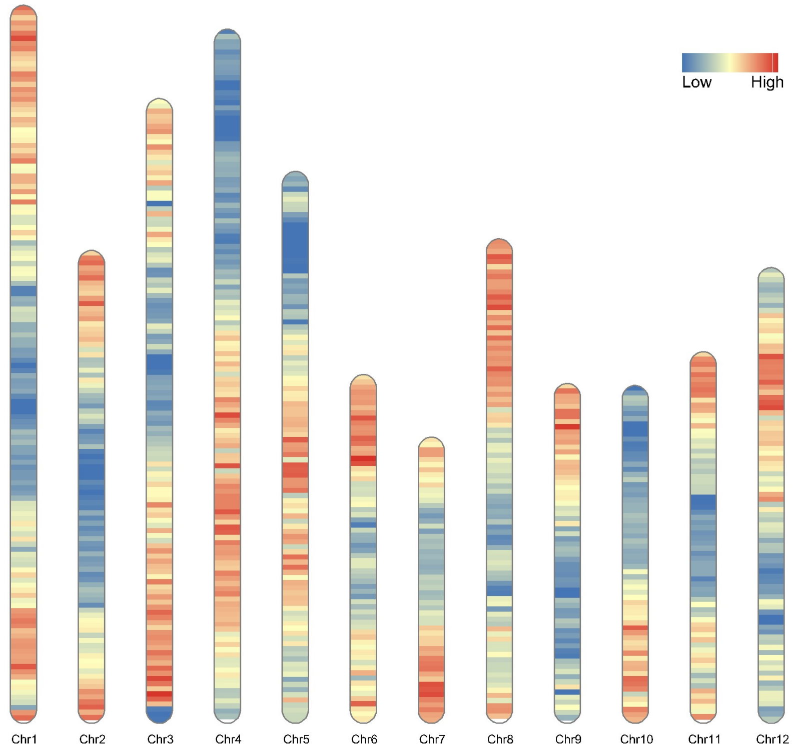


**Supplementary Figure.3 Gene density of 12 chromosomes of *C. burmannii***

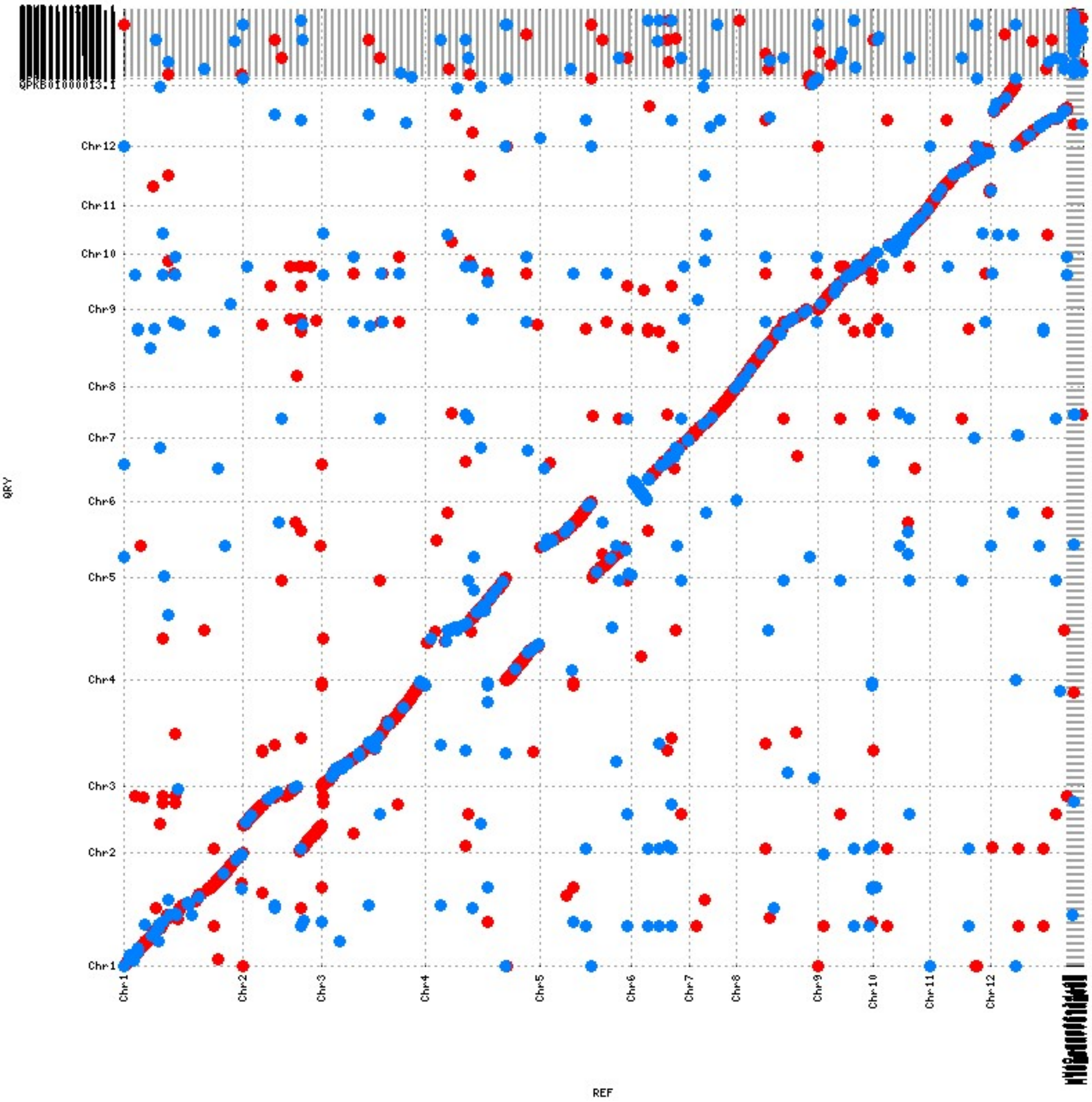


**Supplementary Figure.4 Collinearity of genomic sequences between *C. burmannii* and *C. micranthum***

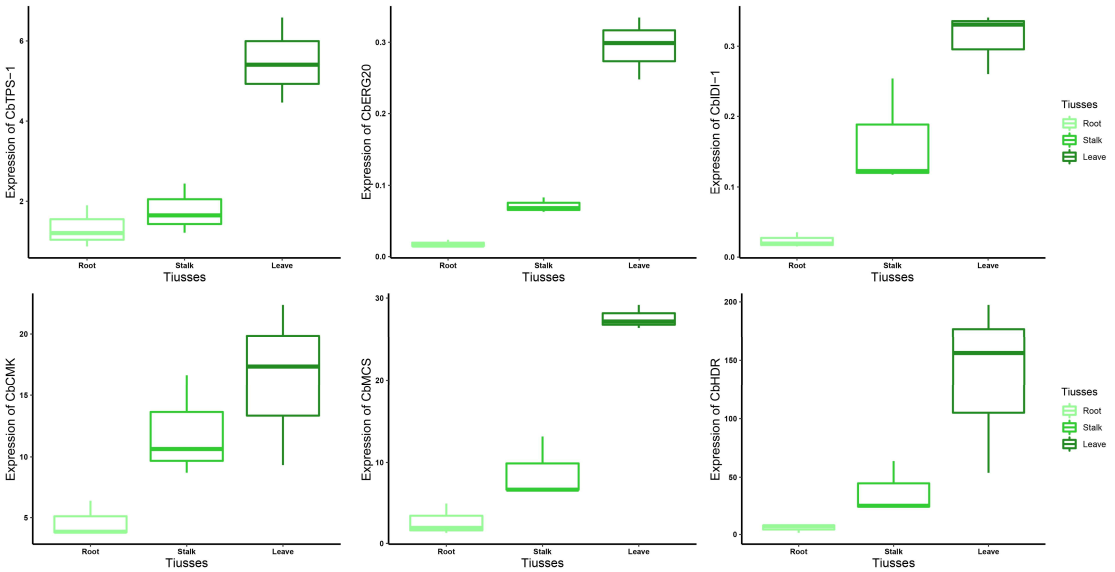


**Supplementary Figure.5**

**qPCR confirmation of partial gene involving in Borneol (+) biosynthesis in *C. burmannii***

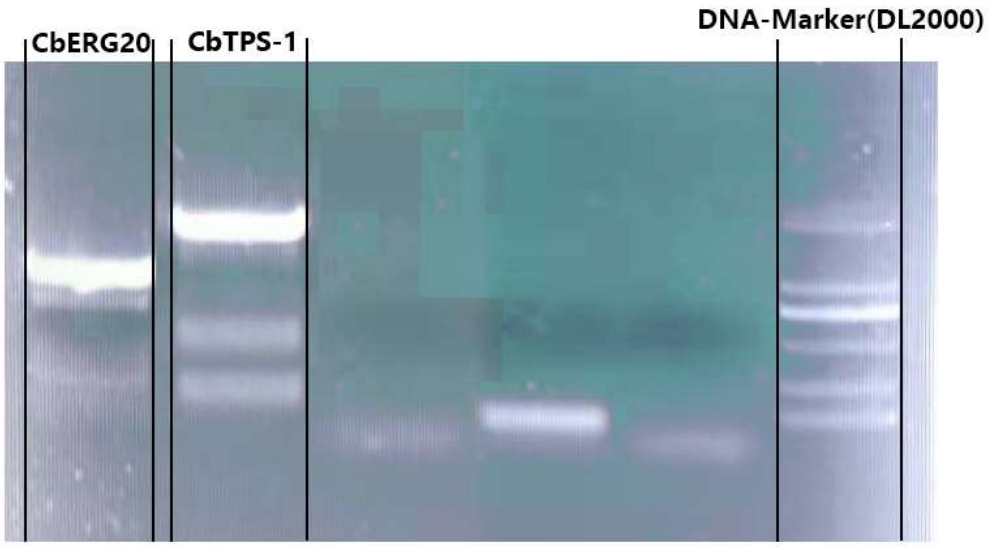


**Supplementary Figure.6**

**Gene clone of *CbERG20* and *CbTPS-1* from *C. burmannii***

**Supplementary Table:**

### Supplementary Table.1

**Genomic survey result of *C. burmannii generated by software Gce***

| Statistics | Value |
|---|---|
| raw_peak | 31 |
| effective_kmer_species | 477904555 |
| effective_kmer_individuals | 36133943881 |
| coverage_depth | 29.7911 |
| genome_size | 1.21E+09 |
| a[1/2] | 0.207058 |
| a[1] | 0.385589 |
| b[1/2] | 0.0676336 |
| b[1] | 0.197718 |
| heterozygosity | 0.68% |

### Supplementary Table.2

**Busco assessment results of *C. micranthum***

| Statistics | Count | Value |
|---|---|---|
| Complete BUSCOs (C) | 2085 | 89.70% |
| Complete and single-copy BUSCOs (S) | 1881 | 80.90% |
| Complete and duplicated BUSCOs (D) | 204 | 8.80% |
| Fragmented BUSCOs (F) | 61 | 2.60% |
| Missing BUSCOs (M) | 180 | 7.70% |
| Total BUSCO groups searched | 2326 | 100% |

### Supplementary Table.3

**Length and types Statistics of LTR-RTs**

| LTR Types (Size/Percents) / species | Copia | Gypsy | Unknown | Total LTR | Difference |
|---|---|---|---|---|---|
| ***C. burmannii*** | 142.94Mb | 244.77Mb | 79.062Mb | 459.42Mb | 272.8Mb |
| | 12.62% | 21.61% | 6.98% | 40.56% | |
| ***C. kanehirae*** | 44.59Mb | 76.05Mb | 65.98Mb | 186.62Mb | |
| | 6.10% | 10.40% | 9.03% | 25.53% | |

### Supplementary Table.4

#### Numbers of TPS subfamilies in the 5 genomes

| TPS subfamilies / Function species | Genome size (Mb) | *TPS-a* | *TPS-b* | *TPS-c* | *TPS-e* | *TPS-f* | *TPS-g* | Total no. |
|---|---|---|---|---|---|---|---|---|
| | | *C15* | *IspS, C10* | *CPS, C20* | *KS, C20* | *C20* | *C10* | |
| ***C. burmannii*** | **1133** | 29 | 23 | 1 | 2 | 6 | 12 | 73 |
| ***C. kanehirae*** | **731** | 28 | 59 | 9 | 3 | 6 | 21 | 126 |
| ***M. acuminata*** | **473** | 26 | 14 | 2 | 0 | 1 | 6 | 49 |
| ***O. sativa japonica*** | **373** | 26 | 0 | 5 | 9 | 0 | 5 | 45 |
| ***D. carota*** | **421** | 1 | 18 | 3 | 2 | 1 | 9 | 34 |
| **Total** | | 110 | 114 | 20 | 16 | 14 | 53 | 327 |

### Supplementary Table.5

#### List about the gene in TPS family or involving in Borneol (+) biosynthesis in *C. burmannii*

| Geneid | Type | Name | Chr | Start | End |
|---|---|---|---|---|---|
| CBB09T444150 | TPS-a | . | Chr9 | 25906084 | 25907964 |
| CBB06T352260 | TPS-a | . | Chr6 | 56816053 | 57117383 |
| CBB09T444170 | TPS-a | . | Chr9 | 25934708 | 25936585 |
| CBB09T444120 | TPS-a | . | Chr9 | 25680221 | 25686069 |
| CBB09T444010 | TPS-a | . | Chr9 | 25488256 | 25494770 |
| CBB09T444260 | TPS-a | . | Chr9 | 26093536 | 26412653 |
| CBB04T249630 | TPS-a | . | Chr4 | 77090824 | 77099858 |
| CBB04T249660 | TPS-a | . | Chr4 | 77156455 | 77165256 |
| CBB09T444310 | TPS-a | . | Chr9 | 26537971 | 26547306 |
| CBB09T443960 | TPS-a | . | Chr9 | 25295398 | 25304493 |
| CBB04T249600 | TPS-a | . | Chr4 | 77023324 | 77059757 |
| CBB03T203250 | TPS-a | . | Chr3 | 76321010 | 76339118 |
| CBB09T444230 | TPS-a | . | Chr9 | 26047759 | 26052339 |
| CBB03T203320 | TPS-a | . | Chr3 | 76402326 | 76406525 |
| CBB03T203390 | TPS-a | . | Chr3 | 76495227 | 76499429 |
| CBB07T359950 | TPS-a | . | Chr7 | 3551214 | 3556335 |
| CBB03T179640 | TPS-a | . | Chr3 | 2847584 | 2851381 |
| CBB09T444300 | TPS-a | . | Chr9 | 26502520 | 26504056 |
| CBB11T099630 | TPS-a | . | Chr11 | 51078989 | 51080137 |
| CBB06T352270 | TPS-a | . | Chr6 | 57124669 | 57128065 |
| CBB11T099660 | TPS-a | . | Chr11 | 51158726 | 51159504 |
| CBB07T359930 | TPS-a | . | Chr7 | 3372526 | 3507321 |
| CBB04T243780 | TPS-a | . | Chr4 | 67877013 | 67881485 |

| | | | | | |
|---|---|---|---|---|---|
| CBB09T444020 | TPS-a | . | Chr9 | 25529991 | 25531006 |
| CBB11T103320 | TPS-a | . | Chr11 | 58939347 | 58939757 |
| CBB11T103310 | TPS-a | . | Chr11 | 58930911 | 58933667 |
| CBB07T359980 | TPS-a | . | Chr7 | 3593445 | 3600025 |
| CBB03T179650 | TPS-a | . | Chr3 | 2862861 | 2863085 |
| CBB09T443930 | TPS-a | . | Chr9 | 25171165 | 25174205 |
| CBB08T411490 | TPS-b | . | Chr8 | 42370152 | 42379403 |
| CBB09T443800 | TPS-b | . | Chr9 | 24800658 | 24811679 |
| CBB09T443810 | TPS-b | . | Chr9 | 24949053 | 24962919 |
| CBB08T411440 | TPS-b | . | Chr8 | 42198126 | 42205994 |
| CBB06T352170 | TPS-b | . | Chr6 | 56640790 | 56657084 |
| CBB06T352490 | TPS-b | . | Chr6 | 57542460 | 57547739 |
| CBB06T352480 | TPS-b | . | Chr6 | 57517330 | 57523606 |
| CBB08T411410 | TPS-b | . | Chr8 | 42147649 | 42156095 |
| CBB06T352510 | TPS-b | . | Chr6 | 57566353 | 57572130 |
| CBB06T352420 | TPS-b | CbTPS-2 | Chr6 | 57404969 | 57436005 |
| CBB06T352630 | TPS-b | . | Chr6 | 57734932 | 57745751 |
| CBB06T352950 | TPS-b | . | Chr6 | 58310159 | 58317187 |
| CBB06T352390 | TPS-b | CbTPS-1 | Chr6 | 57339634 | 57353769 |
| CBB03T216060 | TPS-b | . | Chr3 | 99143303 | 99147570 |
| CBB03T215930 | TPS-b | . | Chr3 | 98849388 | 98852962 |
| CBB03T216050 | TPS-b | . | Chr3 | 99098545 | 99101691 |
| CBB09T443240 | TPS-b | . | Chr9 | 23378971 | 23379249 |
| CBB06T352090 | TPS-b | . | Chr6 | 56350415 | 56350984 |
| CBB06T352160 | TPS-b | . | Chr6 | 56595152 | 56595735 |
| CBB06T352060 | TPS-b | . | Chr6 | 56292515 | 56293094 |
| CBB06T352180 | TPS-b | . | Chr6 | 56680323 | 56680906 |
| CBB06T352070 | TPS-b | . | Chr6 | 56325986 | 56326568 |
| CBB06T351940 | TPS-b | . | Chr6 | 56036743 | 56041760 |
| CBB01T023380 | TPS-c | . | Chr1 | 37744969 | 37891054 |
| CBB06T348880 | TPS-e | . | Chr6 | 50044205 | 50142254 |
| CBB06T347380 | TPS-e | . | Chr6 | 45537879 | 45561501 |
| CBB10T075920 | TPS-f | . | Chr10 | 56666157 | 56675165 |
| CBB10T075800 | TPS-f | . | Chr10 | 56495623 | 56501179 |
| CBB10T075830 | TPS-f | . | Chr10 | 56534152 | 56540322 |
| CBB10T075780 | TPS-f | . | Chr10 | 56384592 | 56398615 |
| CBB06T357800 | TPS-f | . | Chr6 | 67914722 | 67924485 |
| CBB06T357760 | TPS-f | . | Chr6 | 67772342 | 67851664 |
| CBB06T352640 | TPS-g | . | Chr6 | 57763825 | 57768202 |
| CBB06T352540 | TPS-g | . | Chr6 | 57586204 | 57591251 |
| CBB04T258000 | TPS-g | . | Chr4 | 91288339 | 91293293 |
| CBB09T443250 | TPS-g | . | Chr9 | 23386035 | 23386858 |
| CBB03T215890 | TPS-g | . | Chr3 | 98780326 | 98785833 |
| CBB06T348950 | TPS-g | . | Chr6 | 50351833 | 50355339 |

| | | | | | |
|---|---|---|---|---|---|
| CBB05T286130 | TPS-g | . | Chr5 | 25737816 | 25741317 |
| CBB06T348980 | TPS-g | . | Chr6 | 50396447 | 50400185 |
| CBB06T348990 | TPS-g | . | Chr6 | 50402792 | 50407417 |
| CBB09T443770 | TPS-g | . | Chr9 | 24749982 | 24750834 |
| CBB09T443760 | TPS-g | . | Chr9 | 24714388 | 24716338 |
| CBB01T034730 | TPS-g | . | Chr1 | 84233775 | 84238285 |
| CBB01T013640 | MVA-Pathway | CbIDI1 | Chr1 | 20750114 | 20758689 |
| CBB07T368550 | MVA-Pathway | CbERG20 | Chr7 | 29344703 | 29362182 |
| CBB03T211320 | MVA-Pathway | CbERG10 | Chr3 | 91319478 | 91340632 |
| CBB09T440080 | MVA-Pathway | CbERG12 | Chr9 | 17483309 | 17485412 |
| CBB12T110590 | MVA-Pathway | CbERG8-1 | Chr12 | 1862961 | 1883815 |
| CBB12T144310 | MVA-Pathway | CbERG8-2 | Chr12 | 84846922 | 84872759 |
| CBB10T063850 | MVA-Pathway | CbERG19 | Chr10 | 31053941 | 31078924 |
| CBB02T173250 | MVA-Pathway | CbERG13-1 | Chr2 | 85640974 | 85650426 |
| CBB04T273790 | MVA-Pathway | CbERG13-2 | Chr4 | 114421193 | 114431483 |
| CBB08T420550 | MVA-Pathway | CbERG13-3 | Chr8 | 77000979 | 77006911 |
| CBB04T255870 | MVA-Pathway | CbThmg1-1 | Chr4 | 88164670 | 88167281 |
| CBB11T088050 | MVA-Pathway | CbThmg1-2 | Chr11 | 8715487 | 8717338 |
| CBB04T259280 | MVA-Pathway | CbThmg1-3 | Chr4 | 92977882 | 92979768 |
| CBB05T313730 | MVA-Pathway | CbThmg1-4 | Chr5 | 77906138 | 77909488 |
| CBB04T256720 | MVA-Pathway | CbThmg1-5 | Chr4 | 89554009 | 89555872 |
| CBB11T083930 | MVA-Pathway | CbThmg1-6 | Chr11 | 3150028 | 3151759 |
| CBB10T079620 | MVA-Pathway | CbThmg1-7 | Chr10 | 62930767 | 62934949 |
| CBB01T051120 | MEP-Pathway | CbDXS-1 | Chr1 | 126300030 | 126332390 |
| CBB03T180510 | MEP-Pathway | CbDXS-2 | Chr3 | 4393363 | 4401485 |
| CBB03T216610 | MEP-Pathway | CbDXS-3 | Chr3 | 99818706 | 99823813 |
| CBB05T309810 | MEP-Pathway | CbDXS-4 | Chr5 | 71556801 | 71564086 |
| CBB08T405490 | MEP-Pathway | CbDXS-5 | Chr8 | 30030186 | 30038723 |
| CBB09T441360 | MEP-Pathway | CbDXS-6 | Chr9 | 19606881 | 19617174 |
| CBB09T436310 | MEP-Pathway | CbDXR | Chr9 | 11062219 | 11073892 |
| CBB12T119610 | MEP-Pathway | CbYgbP | Chr12 | 20551355 | 20559472 |
| CBB04T240280 | MEP-Pathway | CbYchB | Chr4 | 61009266 | 61017335 |
| CBB06T334970 | MEP-Pathway | CbYgbB | Chr6 | 12431589 | 12433851 |
| CBB10T072420 | MEP-Pathway | CbGcpE-1 | Chr10 | 50865941 | 50874700 |

| | | | | | |
|---|---|---|---|---|---|
| CBB08T422050 | MEP-Pathway | CbGcpE-2 | Chr8 | 80504717 | 80511798 |
| CBB07T378750 | MEP-Pathway | CbLytB | Chr7 | 48786911 | 48792667 |

### Supplementary Table.6

#### Sequence and List of primers in qPCR and gene clone confirmation assays.

| | Geneid | Primer name | Primer Sequence |
|---|---|---|---|
| Qpcr | CBB07T368550 | CbERG20-QF | TCCAAGACCCAGCTTTCGAT |
| | | CbERG20-QR | CGATGCACCAACCAAGAACA |
| | CBB01T013640 | CbIDI1-QF | ATCGCGTTATTGGGCATGAC |
| | | CbIDI1-QR | CAGCAGGTGTTTGTCCAGAC |
| | CBB06T352390 | CbTPS-1QF | GCTTCACACCTTGCCTTTCA |
| | | CbTPS-1QR | TGTCCATGTACCACCTAGCC |
| | CBB04T24028 | CbCMK-QF | CGGCTGGATCAAACTAGCTC |
| | | CbCMK-QR | GCATGAATCCGTTGTTTCAA |
| | CBB06T33497 | CbMCS-QF | TGCCTTTTCGGATAGGACAC |
| | | CbMCS-QR | GCATCCACCACACAATGAAG |
| | CBB07T37875 | CbHDR-QF | TTTGCGATGCTACACAGGAG |
| | | CbHDR-QR | TGCCATAGTGCTCTGCAATC |
| Gene clone | CBB07T368550 | CbERG20-CLF | ATGGCTGCGGCGCCAAATGG |
| | | CbERG20-CLR | CTACTTTTGCCTCTTGTAGATC |
| | CBB06T352390 | CbTPS-1-CLF | ATGGCATTGCAAATGACTAC |
| | | CbTPS-2-CLF | GGCAGATCCACCATCAACATA |
